# Characterizing the soil microbial community associated with the fungal pathogen *Coccidioides immitis*

**DOI:** 10.1101/2024.09.27.615053

**Authors:** Molly Radosevich, Jennifer Head, Lisa Couper, Amanda Gomez-Weaver, Simon Camponuri, Liliam Montoya, John Taylor, Justin Remais

## Abstract

Coccidioidomycosis is a fungal disease affecting humans and other mammals, caused by environmental pathogens of the genus *Coccidioides*. Understanding the ecological factors that shape the distribution of *Coccidioides* in soils is important for minimizing the risk of human exposure, though this remains challenging due to the pathogen’s highly variable spatial distribution. Here, we examined associations between the soil microbial community and *Coccidioides immitis* presence within the Carrizo Plain National Monument—a minimally disturbed grassland ecosystem, and the site of a longitudinal study examining the effects of rodents and their burrows on *C. immitis* presence in soils. Using internal transcribed spacer 2 (ITS2) and 16S sequencing to characterize the soil fungal and bacterial communities, we found over 30 fungal species, including several other members of the Onygenales order, that co-occurred with *Coccidioides* more frequently than expected by chance. *Coccidioides*-positive samples were significantly higher in microbial diversity than negative samples, an association partly driven by higher *Coccidioides* presence within rodent burrows compared to surface soils. Soil source (*i.e.,* rodent burrow versus surface soil) explained the largest amount of variation in bacterial and fungal community diversity and composition, with soils collected from rodent burrows having higher microbial diversity than those collected from adjacent surface soils. While prior evidence is mixed regarding associations between *Coccidioides* and microbial diversity, our study suggests that favorable microhabitats such as rodent burrows can lead to a positive association between soil diversity and *Coccidioides* presence, particularly in otherwise resource-limited natural environments.

## Introduction

*Coccidioides*—a genus of soil-dwelling pathogenic fungi within the *Onygenaceae* family that cause coccidioidomycosis (also known as Valley fever)—is among the priority fungal pathogens of concern identified by the World Health Organization (WHO) (1,2). There is currently no available vaccine to prevent coccidioidomycosis, making reducing exposure to the pathogen the primary method of disease prevention. Yet identifying areas of high risk for human exposure remains challenging, as the pathogen’s presence in the environment varies widely over fine spatial scales, and the drivers of its patchy distribution remain poorly understood (3,4). As such, a better understanding of the ecological factors shaping *Coccidioides*’ distribution in the soil is critical for identifying point sources of pathogen exposure risk to humans.

Both *Coccidioides* species (*i.e.*, *C. immitis*, *C. posadasii*) have a dimorphic life cycle and are found in arid regions of North and South America (5). In the soil environment, *Coccidioides* grows as a network of branching hyphae, potentially obtaining carbon and nutrients from the bodies of dead rodents and other sources of animal keratin shed into burrows and the surrounding soil (5–7). As the hyphae grow, they produce chains of asexual spores known as arthroconidia, which become airborne when the soil is disturbed by excavation or wind erosion (8). When a mammal inhales these spores, the fungus initiates the parasitic phase of its life cycle inside the host lungs, and can disseminate to other parts of the body if the host is unable to control the infection. Upon host death, *Coccidioides* may be released from lung nodules called granulomas inside the host, where it is subsequently hypothesized to utilize the carcass as a source of nutrients (1,5,6).

In regions known to harbor *Coccidioides*, its spatial and temporal distribution in the air and soil are sporadic and uneven (3,4). In some cases, *Coccidioides*-positive sites are identified only in the wake of a coccidioidomycosis outbreak linked to a specific geographic area and timeframe (9–13). However, it is not uncommon, even in studies sampling in putatively positive areas based on epidemiologic data, for the fungus to be detected in few or none of the samples drawn from soils (e.g., <10%; (14). In the air, extensive sampling over regions with consistently high case rates has demonstrated sporadic detection of airborne *Coccidioides*, with a few consistently positive “hot spots”, suggesting a highly uneven distribution and unclear associations between arthroconidia concentrations and environmental conditions such as dust storms (4). Prior limitations in the molecular detection methods for *Coccidioides* may have contributed to the observed unevenness in the soil, but recent advancements in the sensitivity and specificity of PCR assays for the pathogen have minimized this challenge, lowering the limit of detection to as low as <15 target DNA copies per reaction (15).

Many factors are hypothesized to determine the pathogen’s distribution, including seasonal, annual, and interannual climate cycles (16,17); the presence and abundance of specific mammalian host species (6,18,19), and physical and chemical soil properties (7,20). As stated by the endozoan, small-mammal reservoir hypothesis, *Coccidioides* spp. are posited to persist in the soil primarily through infection of wild populations of rodents and other small mammals (6). Several studies have found that *Coccidioides* is more often detected inside rodent burrows than in the surrounding soil (14,19,21). However, the relative roles of rodents themselves versus the burrows they create in structuring where *Coccidioides* spp. are found in the environment remain unclear. Recent work found that rodent burrow creation mediates the effect of rodents on *Coccidioides* presence, with 73.7% of the association between rodents and *Coccidioides* attributable to the creation of burrows (21). Our current research is motivated by understanding how the relationships between rodents, burrows, and *Coccidioides* presence may be impacted by soil microbial community dynamics. Determining how the soil microbial community may influence *Coccidioides*’ ecology and structure *Coccidioides* populations in the soil will inform recommendations on minimizing human exposure risk to the pathogen.

Soil microbial community dynamics may play a role in determining where *Coccidioides* is found in the soil. As antifungal activity is relatively common amongst soil bacteria (22,23), some studies have explored the potential role of soil microbial community composition and diversity and the presence of specific fungal or bacterial taxa in determining *Coccidioides* presence in the environment and growth on plates (7,24–27). For instance, researchers characterized the fungal community of soil taken from a known *Coccidioides immitis*-positive site in eastern Washington state, finding qualitative differences in the most abundant taxa across taxonomic levels between *Coccidioides*-positive and *Coccidioides*-negative samples (7). Additionally, *Coccidioides immitis* presence was positively associated with the genus *Aureobasidium* (containing soil saprotrophs), and negatively associated with the genus *Phialocephala* (containing plant endophytes), aligning with evidence that *Coccidioides* was primarily found at sites lacking vegetation (7). In the southern San Joaquin Valley, California— the region in which the vast majority of coccidioidomycosis cases are reported in the state each year—prior research found no strong association between the presence of *Coccidioides* and the overall fungal community composition, though 19 fungi were identified as potential indicator species (16 positively, 3 negatively associated with *Coccidioides*) (24). Other studies have found that naturally occurring microbes isolated from the soil at *Coccidioides*-positive sites exhibit antifungal activity toward the pathogen, inhibiting its growth on plates, suggesting that *Coccidioides* is a poor competitor against other soil microorganisms (25,26,28).

While individual fungal and bacterial taxa have been investigated as potential microbial antagonists to *Coccidioides*, their impact in natural, ecological settings is unclear. That is, the relationship between natural soil bacterial and fungal communities and *Coccidioides* presence, and whether and how this relationship may be mediated by the soil microhabitat, remains poorly understood. Herein, we leveraged a long-term ecological experiment in the Carrizo Plain National Monument in California—a region known to harbor *Coccidioides immitis* (29)—to investigate the relationship between soil microbial community dynamics and *C. immitis* presence across varying relevant microhabitats. Specifically, we collected soil samples from a series of replicated experimental plots and analyzed the soil microbial communities from inside active rodent burrows—known to harbor *Coccidioides* at greater frequency (19,21)—and from burrows from which rodents had been excluded for the previous 14 years. We additionally collected soils from the ground surface nearby both active and inactive burrows. This design allowed us to disentangle the influence of rodent presence and the burrow microhabitat on soil microbial dynamics and *Coccidioides* presence—a key open question (21). Finally, we investigated whether specific fungal or bacterial taxa have positive or negative co-occurrence patterns with *C. immitis*. Our primary findings that *Coccidioides*-positive soils contain higher microbial diversity than negative soils, and that *Coccidioides* frequently co-occurs with phylogenetically similar fungi, suggest that these organisms may compete for similar resources in the burrow microenvironment. Our study highlights the importance of continued environmental sampling and investigation of *Coccidioides* in its natural environment and has implications for identifying sources of *Coccidioides* in the soil and modeling the distribution of the pathogen.

## Methods

### Soil Collection

We collected soil samples from study sites within the Carrizo Plain National Monument in eastern San Luis Obispo County, California (Figure 1A), an ecosystem characterized by extensive rodent burrow systems where an ongoing, long-term experiment is examining the impacts of drought, livestock management, *Dipodomys ingens* (giant kangaroo rat) activity and the presence of other wildlife on ecological communities (30,31). Samples were collected across two pastures (hereafter, “sites”) in the monument. Samples from both sites were included in analyses to increase replication of our findings across different locations. The northern site periodically experiences seasonal cattle grazing and is dominated by several exotic annual grasses, while the southern site has never been grazed and is a dominated by a single native perennial grass species (30). Within each site, we collected samples from 8 plots, each measuring 140 x 140 meters and containing a 20 x 20 meter rodent exclosure fence established in 2007 at the center (30) (Figure 1B). The rodent exclosure fence extends 61 cm into the soil and 91 cm above the ground surface. At the time of establishment, and thereafter, monthly, any rodents found inside the fence were relocated elsewhere (30). Outside of the rodent exclosure, where there is active rodent presence, we randomly selected one rodent burrow system (a precinct) and collected replicate precinct samples from inside the mouth of five burrows; these samples were paired with five replicate topsoil samples collected starting at two meters away from the nearest burrow entrance, and subsequently at one-meter intervals, from a depth of 10 centimeters below the ground surface. Additionally, we sampled five replicate precinct (abandoned since 2007) and topsoil samples from inside the exclosure following the same procedure. All samples analyzed were collected in April 2021.

**Figure 1.**
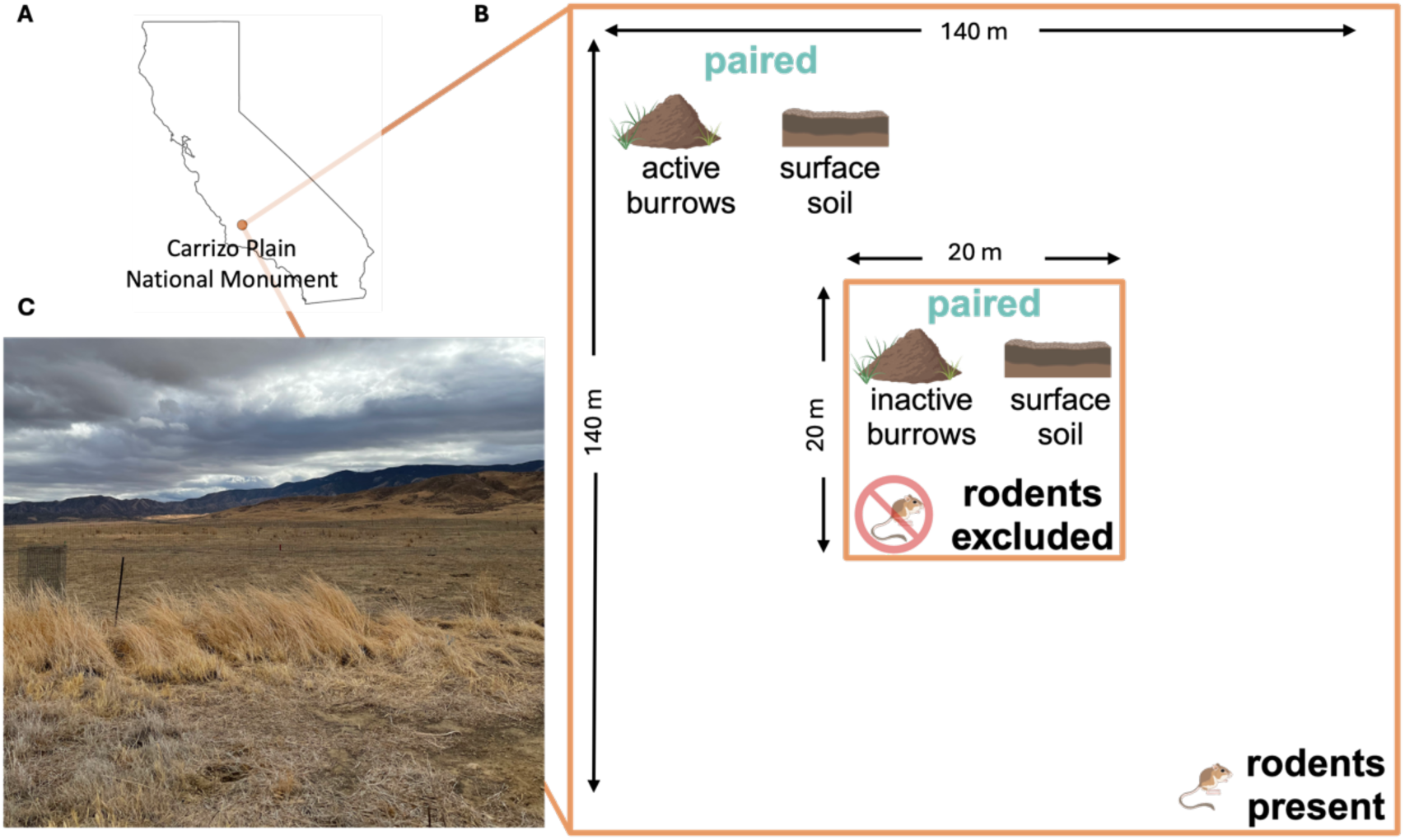
(A) Location of the Carrizo Plain National Monument (orange point) in California. (B) Schematic of one sampling plot, comprising an inner zone (center; 20 m x 20 m) where rodents are excluded by a physical barrier (exclosure), surrounded by an outer zone (140 m x 140 m) where rodents are present (Note: figure not drawn to scale). Five replicate precinct soil samples are collected from inactive burrows within the rodent exclosure and paired with five adjacent soil samples collected from the ground surface within the exclosure (see Methods: *Soil Collection*). Paired precinct samples and adjacent surface soil samples are also collected in the surrounding zone where rodents are present. (C) A photograph from the field site, a grassland plain bordered to the northeast by the Temblor Range and the southwest by the Caliente Range, taken in October 2021.

### Soil DNA Extraction

We extracted genomic DNA from all soil samples using the DNeasy PowerSoil Pro Kit (QIAGEN, Hilden, Germany), with modifications to the protocol in accordance with Biosafety Level 2+ control measures. For each sample, we transferred approximately 200-500 µL of soil into a 2-mL Lysing Matrix E tube prefilled with a mixture of ceramic, silica, and glass beads (MPBio, Burlingame, California, USA) and containing 800 µL of lysis buffer (Buffer CD1). Samples were homogenized in an MP Bio FastPrep-24 5G Cell Disruptor for 2 cycles of 30 seconds each at 6 m/s, separated by a five-minute rest period. Then, we centrifuged samples at 10,000 x g for 1 minute and transferred the supernatant to a clean 2-mL tube. Past this point, we conducted the remainder of the protocol as specified by the DNeasy PowerSoil Pro Kit manual on the laboratory benchtop.

### qPCR to Detect Coccidioides spp

After DNA extraction and quantification via the Qubit^TM^ dsDNA Quantification High Sensitivity Assay Kit (ThermoFisher Scientific, Waltham, Massachusetts, USA), we diluted samples to 12 ng/µL in preparation for qPCR (*i.e.,* to minimize PCR inhibition occurring at higher DNA concentrations as in Bowers et al. (2019) (15). We conducted qPCR following the CocciENV assay, and processed all samples in quadruplicate plate wells (15,32). Samples were considered positive for *Coccidioides* spp. if at least 3 out of 4 wells had a Ct value below 40 (15,24).

### ITS2 and 16S Amplicon Sequencing

After assessing the presence of *Coccidioides* in each sample, we prepared samples for PCR amplification and pooling into both internal transcribed spacer 2 (ITS2) libraries (fungi) and 16S libraries (bacteria). ITS2 is a highly conserved region of nuclear ribosomal DNA that allows for discrimination between species of fungi (33–35). Similarly, the 16S gene region encodes for ribosomal RNA and is commonly used to discriminate between genera or families of bacteria (36). We used 5.8S-Fun and ITS4-Fun primers for ITS2 sequence amplification (37,38), and 341F and 785R primers for amplification of the 16S V3-V4 gene region (39). We set a target sample size of 300 samples to obtain sufficient sequencing depth (averaging 100,000 reads per sample). Libraries were sent to the Vincent J. Coates Genomic Sequencing Laboratory (QB3 Genomics, UC Berkeley, Berkeley, CA, RRID:SCR_022170) for fragment analysis and sequencing using the Illumina MiSeq v3 300 PE kit (Illumina, Inc., San Diego, CA, USA). The PCR amplification protocol was based on methodology detailed elsewhere (38,39).

### Amplicon Sequence Processing

After sequencing and demultiplexing, we processed the sequences through the QIIME2 pipeline (40). We used Cutadapt to trim primers out of all sequences (41), and used a minimum quality score cutoff of 25 to identify where to truncate all sequences before downstream analysis. We then processed the sequences in DADA2 to truncate and merge forward and reverse sequences, using the consensus method to exclude chimeras (42). Finally, all merged amplicon sequence variants (ASVs) were assigned taxonomy using the UNITE database for fungi (97% similarity) (43) and the SILVA database for bacteria (99% similarity) (44).

### Statistical Analyses

Using fungal and bacterial DNA sequences, we estimated microbial community diversity and composition via the Shannon index and Bray-Curtis dissimilarity to estimate alpha and beta diversity, respectively (45,46). We also estimated co-occurrence patterns with *Coccidioides* via checkerboard score analysis (47). For all analyses, we analyzed ITS2 and 16S data separately, assessing ITS2 data at the species level and 16S data at the family level. We treated the following variables as binary for all analyses: sample source (burrow vs. surface); rodent exclosure status (rodents present vs. excluded); *Coccidioides* status (positive vs. negative); and site of collection (southern vs. northern pasture).

We estimated alpha diversity (richness, evenness, and Shannon index) for fungal and bacterial sequences using the ‘phyloseq’ package in R (45,48), and compared mean alpha diversity measures across *Coccidioides* status, sample source, rodent exclosure status, and site using the Wilcoxon rank-sum test. Additionally, leveraging the paired burrow-surface soil sampling design at each plot, we conducted a Wilcoxon signed-rank test to directly compare the average alpha diversity of replicate precinct samples to that of adjacent replicate surface soil samples, both within and outside of rodent exclosures.

We characterized beta diversity, a measure of the pairwise differences in microbial community composition between soil samples, separately for fungi and bacteria. We visualized differences in beta diversity based on sample source, rodent exclosure status, and *Coccidioides* status using Principal Coordinate Analysis (PCoA) and analyzed the differences using a nested non-parametric permutational multivariate analysis of variance (PerMANOVA) test. First, we transformed ASV sequence data using square-root and Wisconsin double-standardization, and then generated a Bray-Curtis dissimilarity matrix using the vegan package (46,49). We used dimension reduction to visualize the dissimilarity matrices via generating the top two principal coordinates explaining the most variation between the samples. Finally, we conducted a nested PerMANOVA to see whether microbial community composition differed significantly based on any experimental variables in the dataset (50), adjusting for multiple comparisons using false discovery rate correction (51).

We used checkerboard score analysis, in which counts of pairwise taxa co-occurrences are compared to null expectations based on permuting taxa counts across samples, to examine pairwise taxa associations with *Coccidioides*, using the R package ‘ecospat’ version 4.0.0 (47,52). This analysis is used to determine whether two taxa occur together in a sample more or less often than would be expected by chance. The observed checkerboard scores (C-scores) for each pair of taxa were generated from a presence/absence sample-by-taxa data matrix. We conducted 1000 permutations under a fixed-equiprobable null model, where column (taxa) sums are fixed but sample labels are randomly shuffled, generating a distribution of expected C-scores under the null hypothesis. Finally, we compared the mean expected C-score to the observed C-score to generate a standardized effect score (SES) for each pair of taxa, as well as a significance level, which was adjusted (false discovery rate correction) to account for the number of comparisons (51). A positive SES indicates that two species co-occurred less often than expected under the null, and a negative SES indicates that they occurred more often than expected. All statistical analyses were conducted in R version 4.3.0.

## Results

We analyzed a total of 318 soil samples (Table S1) via 16S and ITS2 sequencing to investigate the microbial community and its relationship to *Coccidioides immitis* presence. Based on qPCR analysis, 11.9% (38/318) of all soil samples were positive for *Coccidioides* spp. with substantially higher positivity rates for samples from rodent burrows (21.4%, 34/159) compared to surface soils (2.5%, 4/159). Within the *Coccidioides*-positive rodent burrow samples, 67.6% (23/34) were from burrows with active rodent presence, while 32.3% (11/34) were from inactive burrows.

We obtained a total of 24,053,261 reads from ITS2 sequencing and 25,709,195 from 16S sequencing, with 532 ASVs matched to known fungal species and 394 ASVs matched to known bacterial families. Within the known taxa, ITS2 sequencing returned a total of 9,731,837 reads (mean number of reads per sample 30,603), while 16S sequencing returned a total of 12,197,453 reads (mean number of reads per sample 38,357). Rarefaction curves indicated that all samples obtained sufficient sequencing depth (Figure S1).

Overall, 65.98% of the unique fungal species in the dataset belonged to the phylum Ascomycota, 22.74% to Basidiomycota, 4.14% to Mucoromycota, and the remainder to Chytridiomycota, Glomeromycota, Mortierellomycota, and Olpidiomycota. *Coccidioides*- positive samples generally had a higher proportion of Ascomycota than *Coccidioides*-negative samples did, and a lower proportion of Basidiomycota (Figure S2). For bacteria, 39 phyla were represented, and the three phyla containing the most taxa were Actinobacteriota, Bacteroidota, and Proteobacteria (Figure S3).

### Alpha Diversity

Across all samples, mean fungal species richness and overall fungal diversity (Shannon index) were significantly higher in *Coccidioides*-positive soils (richness = 81.58) compared to negative soils (richness = 63.70, p < 0.0001) (Table 1, Figure 2A). Additionally, mean bacterial family richness, evenness, and diversity were significantly higher in *Coccidioides*-positive (richness = 141.68) compared to negative soils (richness = 136.50, p < 0.05) (Table 2, Figure 2C). Since most *Coccidioides*-positive samples were also burrow-associated, we further analyzed only soil samples from burrows and found that mean fungal species richness was significantly higher in the *Coccidioides*-positive burrow samples (83.88) than the negative burrow samples (75.06, p < 0.01; Figure 3A, Table 1). This difference was driven by burrow samples from where rodents were excluded. In the burrow samples from where rodents were excluded, fungal species richness was significantly higher in samples with *Coccidioide*s (83.1) versus without (69.6, p-value < 0.01; Figure 3C). Conversely, in the burrow samples from where there was active rodent presence, there was no difference in fungal species richness between samples with or without *Coccidioides* presence. (Figure 3B). Similar trends were observed for bacterial diversity, with *Coccidioides*-positive burrow samples having higher taxa richness than the *Coccidioides*- negative soils, although this was additionally observed when stratifying by burrows with rodent presence and exclusion. However, these associations were non-significant (Supplemental Figure S4).

**Figure 2.**
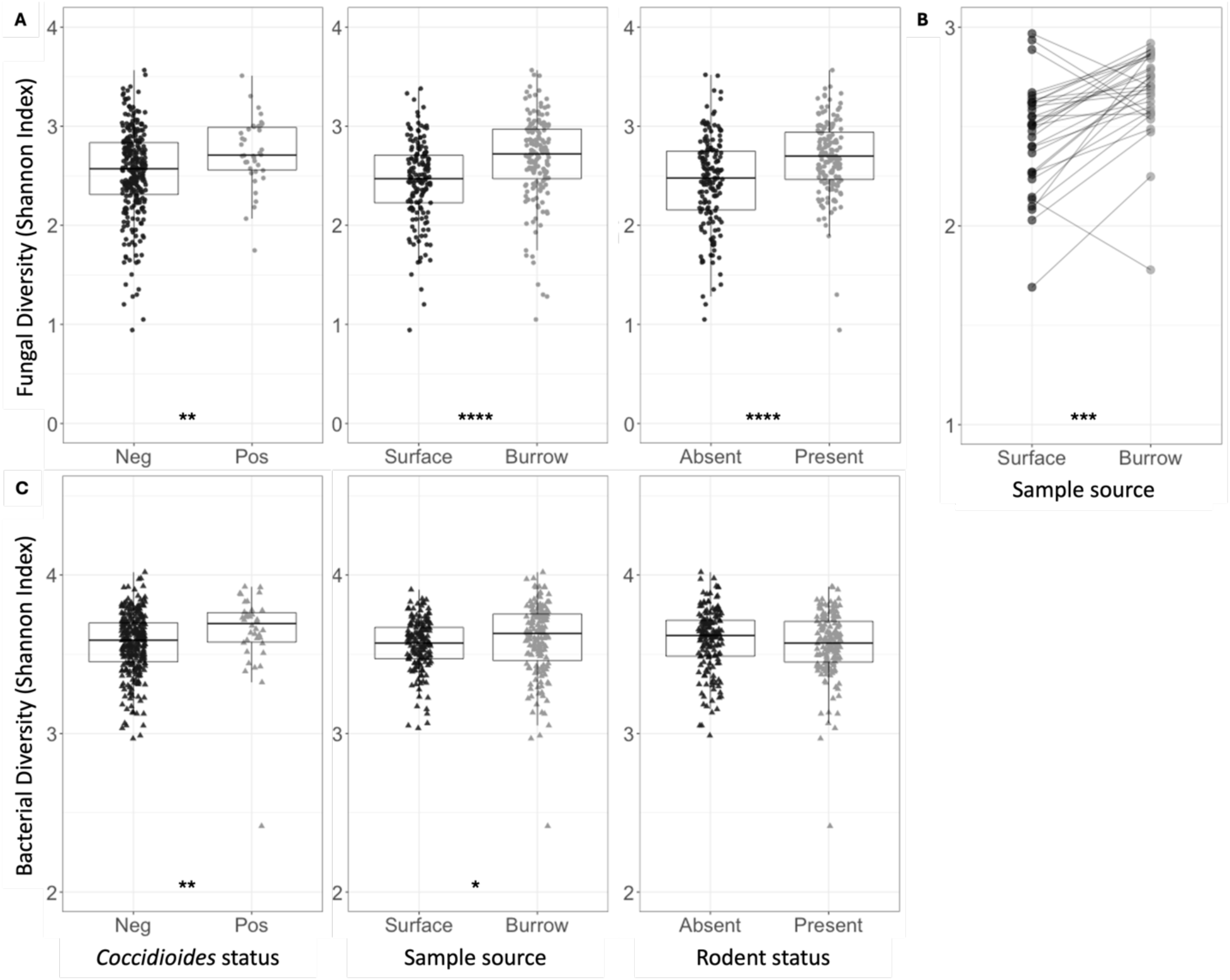
(A) Shannon indices for fungal species diversity of all soil samples, grouped and colored based on *Coccidioides* status, surface versus burrow status (regardless of rodent presence), or rodent status. Each point represents one soil sample. (B) Shannon indices for fungal species diversity of soil samples, averaged across 5 replicate samples collected at each plot. Average estimates for surface soil samples are plotted on the left and burrow soil samples on the right. Lines are drawn to connect spatially paired samples to each other. (C) Shannon indices for bacterial family diversity of all soil samples, grouped and colored based on *Coccidioides* status, surface versus burrow status (regardless of rodent presence) or rodent status. Stars indicate degree of significance based on a Wilcox test, such that * = P ≤0.05, ** = P ≤0.01, *** = P ≤0.001, and **** = P ≤0.0001. No stars indicate P > 0.05.

**Figure 3.**
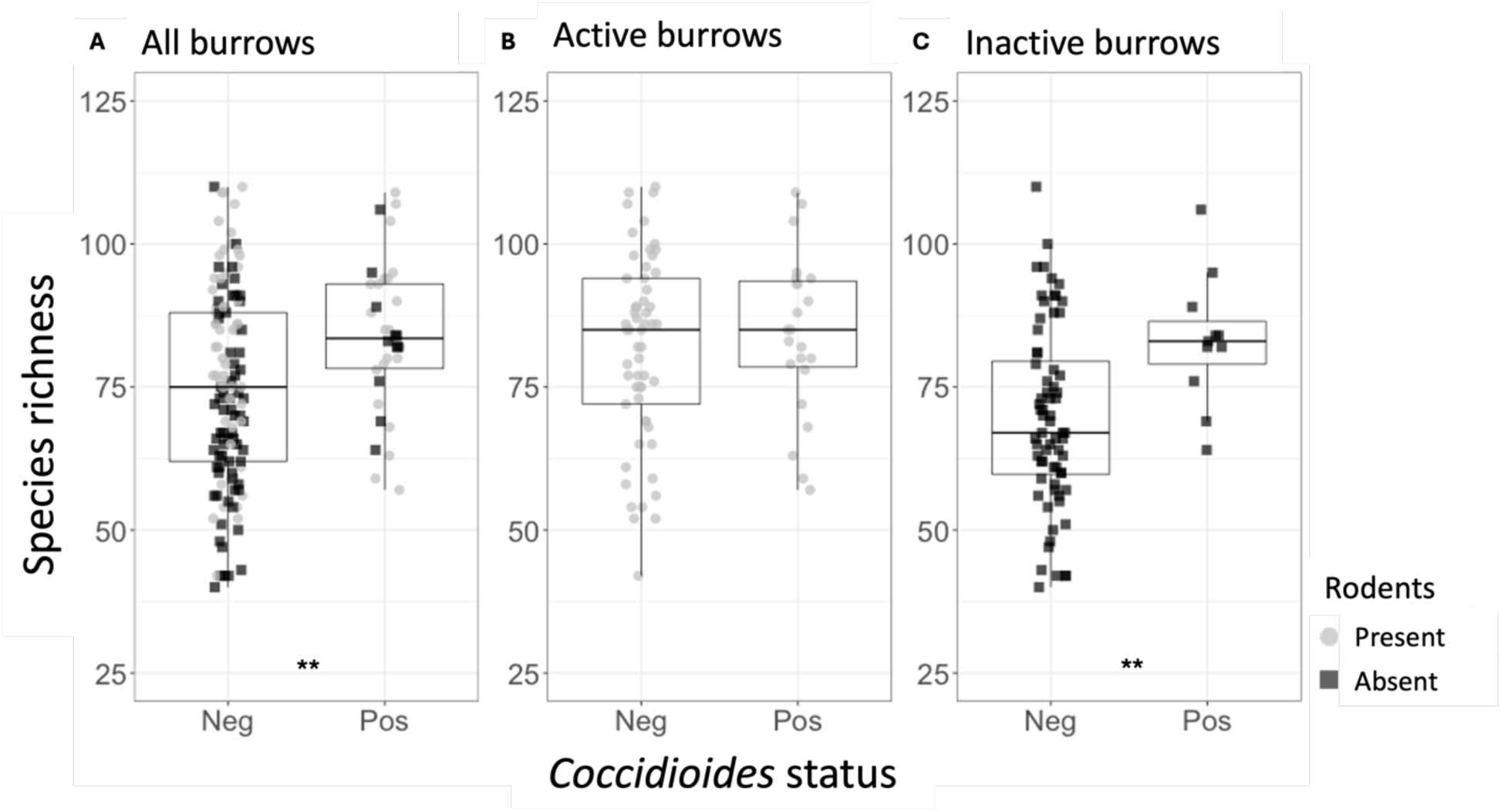
(A) Fungal species richness for the rodent burrow samples, separated by *Coccidioides* status (Neg = *Coccidioides*-negative, Pos = *Coccidioides*-positive). Points represent individual samples and are color-coded based on whether they were taken from active (gray) or inactive (black) rodent burrows. (B) Fungal species richness for the active rodent burrow samples, separated by *Coccidioides* status. (C) Fungal species richness for the inactive rodent burrow samples, separated by *Coccidioides* status. Stars indicate degree of significance based on a Wilcox test, such that * = P ≤0.05, ** = P ≤0.01, *** = P ≤0.001, and **** = P ≤0.0001. No stars indicate P > 0.05.

**Table 1.** Mean fungal species alpha diversity measures (richness, evenness, and Shannon index) for all soil samples. Overall diversity metrics for the whole sample set are reported, as are metrics for subsets of samples grouped based on the following measured variables: pasture, sample source, rodent exclosure status, and *Coccidioides* status. Additionally, alpha diversity metrics for all burrow samples are reported, stratified based on *Coccidioides* status and further based on rodent exclosure status.

|  | Richness<br>Mean (Q1, Q3) | Evenness<br>Mean (Q1, Q3) | Shannon<br>Mean (Q1, Q3) |
| --- | --- | --- | --- |
| **Overall (n = 318)** | 65.84 (51.25, 80.0) | 0.62 (0.57, 0.67) | 2.56 (2.34, 2.85) |
| Northern site (n = 158) | 67.32 (54.00, 83.25) | 0.60 (0.56, 0.66) | 2.53 (2.34, 2.81) |
| Southern site (n = 160) | 64.38 (49.75, 79) | 0.63 (0.58, 0.68) | 2.59 (2.33, 2.88) |
| Burrow (n = 159) | 76.94 (64.50, 89.00) | 0.62 (0.58, 0.67) | 2.67 (2.47, 2.97) |
| Surface (n = 159) | 54.74 (46.00, 61.50) | 0.61 (0.56, 0.67) | 2.44 (2.23, 2.71) |
| Rodents absent (n = 159) | 61.26 (48.50, 71.00) | 0.59 (0.54, 0.66) | 2.44 (2.16, 2.75) |
| Rodents present (n = 159) | 70.42 (54.00, 86.00) | 0.64 (0.60, 0.68) | 2.68 (2.46, 2.94) |
| *Coccidioides*-positive (n = 38) | 81.58 (69.75, 92.25) | 0.62 (0.58, 0.67) | 2.73 (2.56, 2.99) |
| *Coccidioides*-negative (n = 280) | 63.70 (50.00, 75.25) | 0.61 (0.57, 0.67) | 2.54 (2.31, 2.83) |
| **Burrow (n = 159)** |  |  |  |
| *Coccidioides*-positive (n = 34) | 83.88 (78.25, 93.00) | 0.62 (0.57, 0.67) | 2.73 (2.54, 2.99) |
| Rodents absent (n = 11) | 83.09 (79.00, 86.50) | 0.62 (0.57, 0.68) | 2.75 (2.60, 2.98) |
| Rodents present (n = 23) | 84.26 (78.50, 93.50) | 0.62 (0.58, 0.67) | 2.72 (2.53, 2.98) |
| *Coccidioides*-negative (n = 125) | 75.06 (62.00, 88.00) | 0.62 (0.58, 0.67) | 2.66 (2.42, 2.96) |
| Rodents absent (n = 68) | 69.60 (59.75, 79.50) | 0.61 (0.57, 0.66) | 2.57 (2.38, 2.85) |
| Rodents present (n = 57) | 81.56 (72.00, 94.00) | 0.63 (0.60, 0.68) | 2.77 (2.56, 3.02) |

**Table 2.**
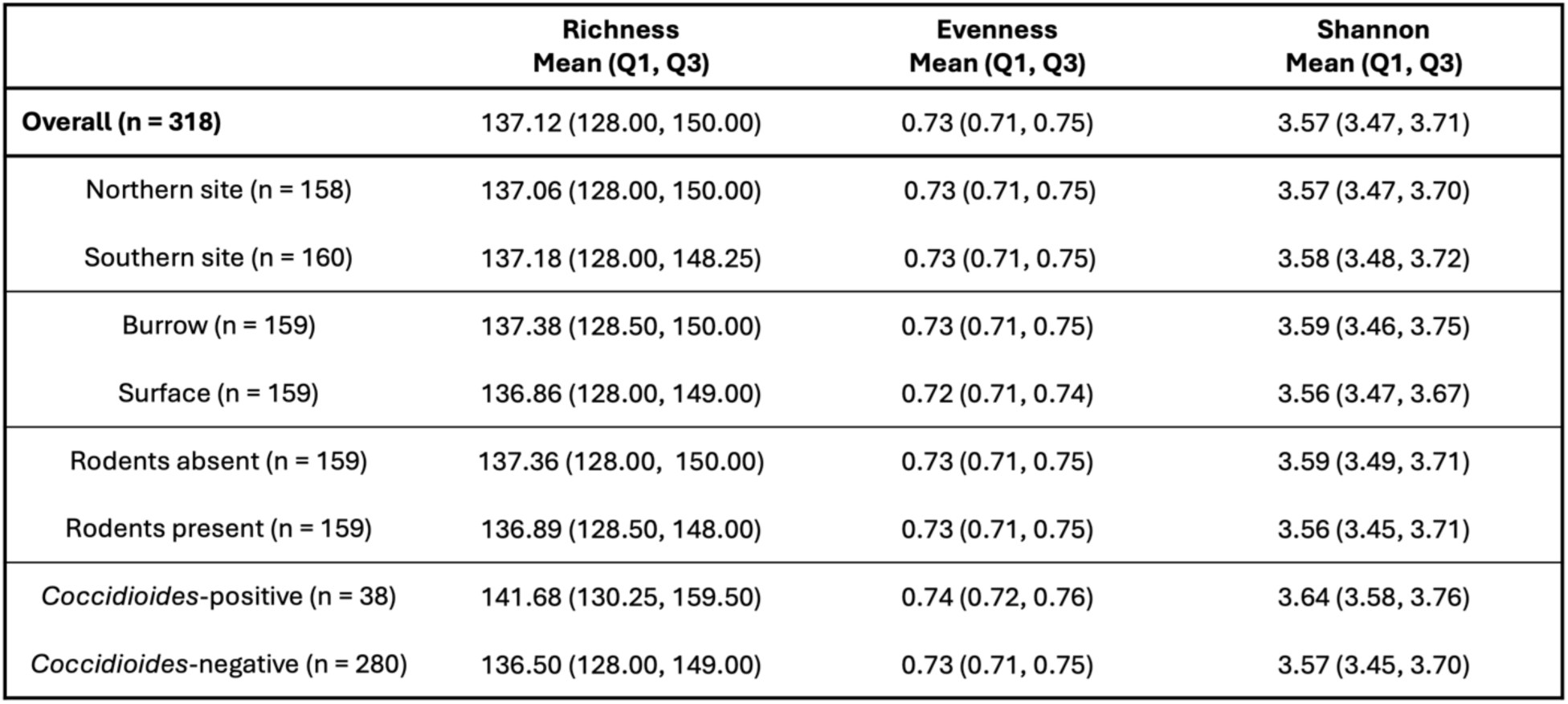
Mean bacterial family alpha diversity measures (richness, evenness, and Shannon index) for all soil samples. Overall diversity metrics for the whole sample set are reported, as are metrics for subsets of samples grouped based on the following measured variables: Pasture of collection, sample source, rodent exclosure status, and *Coccidioides* status.

Mean fungal species richness was significantly higher in rodent burrow soils (76.94) compared to surface soils (54.74, p < 0.0001), and in soils with active rodent presence (70.42) compared to rodent exclosures (61.26, p < 0.0001) (Table 1). Fungal species richness was significantly higher in burrow samples compared to surface samples both within (71.48 vs. 51.18, p < 0.0001) and outside (82.34 vs. 58.34, p < 0.0001) of rodent exclosures. These patterns were recapitulated for fungal species diversity (Shannon index) (Figure 2A, Figure S5, Table 1). Fungal species evenness was significantly higher in soils with rodents present (0.64) versus excluded (0.59, p < 0.0001). Comparing paired burrow-surface soil samples at each plot (Figure 1B), fungal species richness and overall alpha diversity (Shannon index) were higher in burrow soils compared to paired surface soils (p < 0.001 for Shannon index) (Figure 2B). There were no differences in fungal species evenness across paired samples and no trends in bacterial family diversity metrics between paired burrow and surface soils. Despite differences in soil type and cattle grazing, microbial diversity was similar across both sampling sites (Table 1 and Table 2).

Average bacterial family richness did not vary based on other variables aside from *Coccidioides* status (*i.e.,* rodent exclosure status, burrow vs. surface soils) (Table 2). Bacterial diversity and evenness were higher in burrow soils versus surface soils (Figure 2C). Within the rodent exclosure, bacterial evenness and diversity were significantly higher in burrow samples (Shannon index: 3.63) compared to surface soil samples (3.55, p < 0.001), but the same pattern was not observed outside the exclosure. Bacterial diversity metrics did not vary between samples where rodents were present versus excluded (Figure 2C).

### Beta Diversity

Both fungal and bacterial beta diversity varied significantly among samples based on all individual variables, including *Coccidioides* status, sample source, and exclosure status, and site. Of these factors, sample source (burrow vs. surface) explained the largest proportion of variation in microbial community composition (for fungi R^2^ = 7%; for bacteria R^2^ = 9%), followed by site (for fungi R^2^ = 3%; for bacteria R^2^ = 2%), exclosure status (for fungi R^2^ = 3%; for bacteria R^2^ = 1%) and *Coccidioides* status (for fungi R^2^ = 1%; for bacteria R^2^ = 1%; all p < 0.01) (Figure 4). Interactions between variables explained an additional 4% of variation for fungi and 3% of variation for bacteria. Visualization via principal coordinates analysis showed that samples cluster distinctly based on sample source, and, within the fungal dataset only, *Coccidioides* status (Figure 4A).

**Figure 4.**
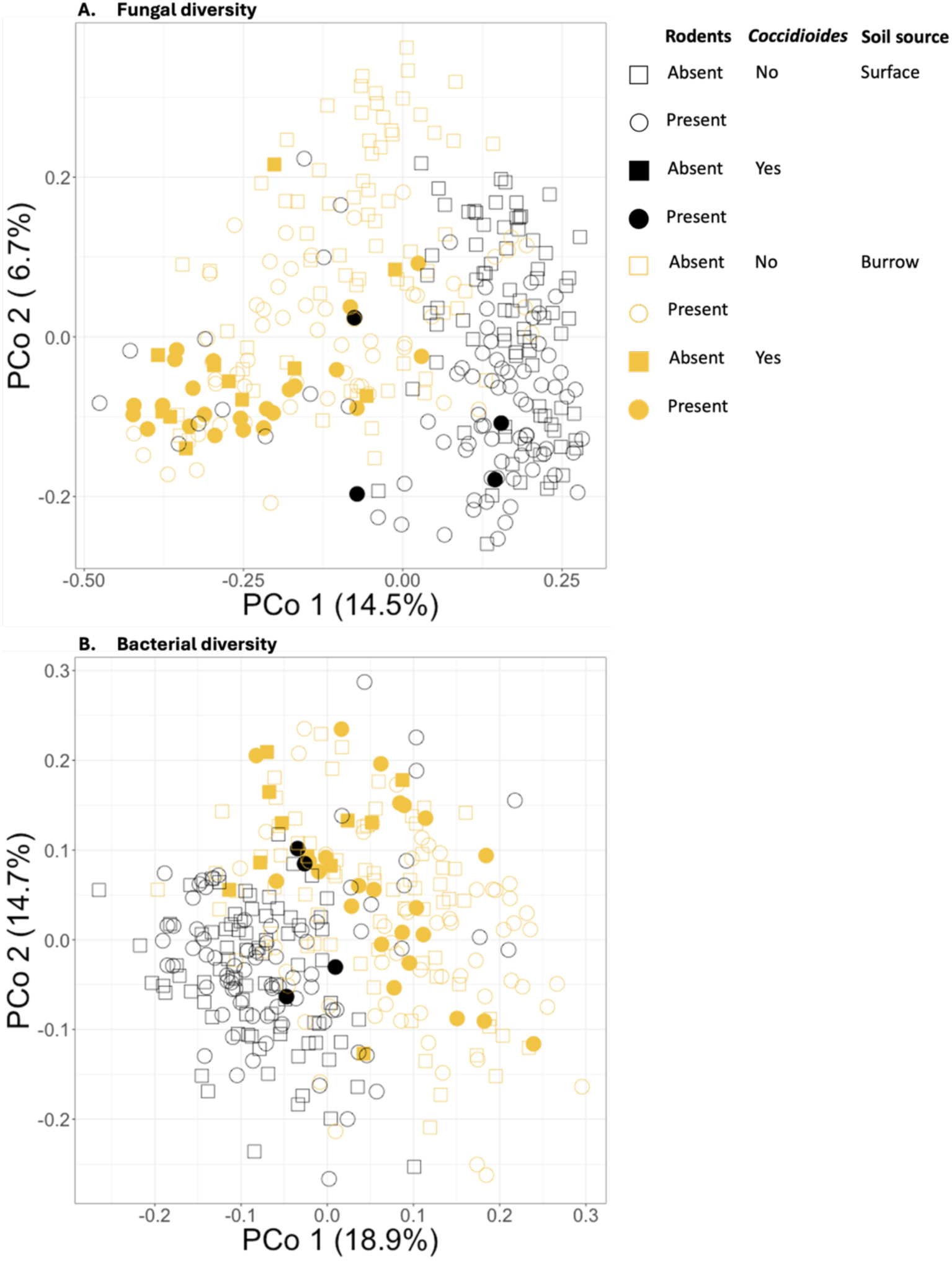
(A) Principal coordinates analysis (PCoA) plot showing Bray-Curtis dissimilarities for the soil fungal communities of all samples. (B) PCoA plot showing Bray-Curtis dissimilarities for the soil bacterial communities of all samples. Samples drawn from burrows are plotted in orange and those from the ground surface in black. Circles represent samples taken from areas with rodent presence and squares represent samples taken from rodent exclosures*. Coccidioides*- positive samples are represented by filled shapes, while negative samples are empty shapes.

### Co-occurrence Analysis

Based on the checkerboard analysis, 37 species of fungi (6.95% of the full fungal dataset) co-occurred with *Coccidioides* significantly more often than expected by chance (Figure 5). No fungal species in our dataset co-occurred with *Coccidioides* less frequently than expected, and no bacterial families had a significant co-occurrence pattern with *Coccidioides* in either direction. Members of the class Eurotiomycetes, order Onygenales, and family *Onygenaceae*, to which *Coccidioides* spp. belongs, were all overrepresented in the set of significantly co-occurring fungal taxa compared to the full dataset (Eurotiomycetes: 37.84% vs. 15.23% in the full dataset; Onygenales: 18.92% vs. 6.2%; and *Onygenaceae*: 2.7% vs. 1.69%). Notably, one species that was identified via our co-occurrence analysis, *Aspergillus penicillioides*, was identified in another study as a potential indicator species for *Coccidioides* (24).

**Figure 5.**
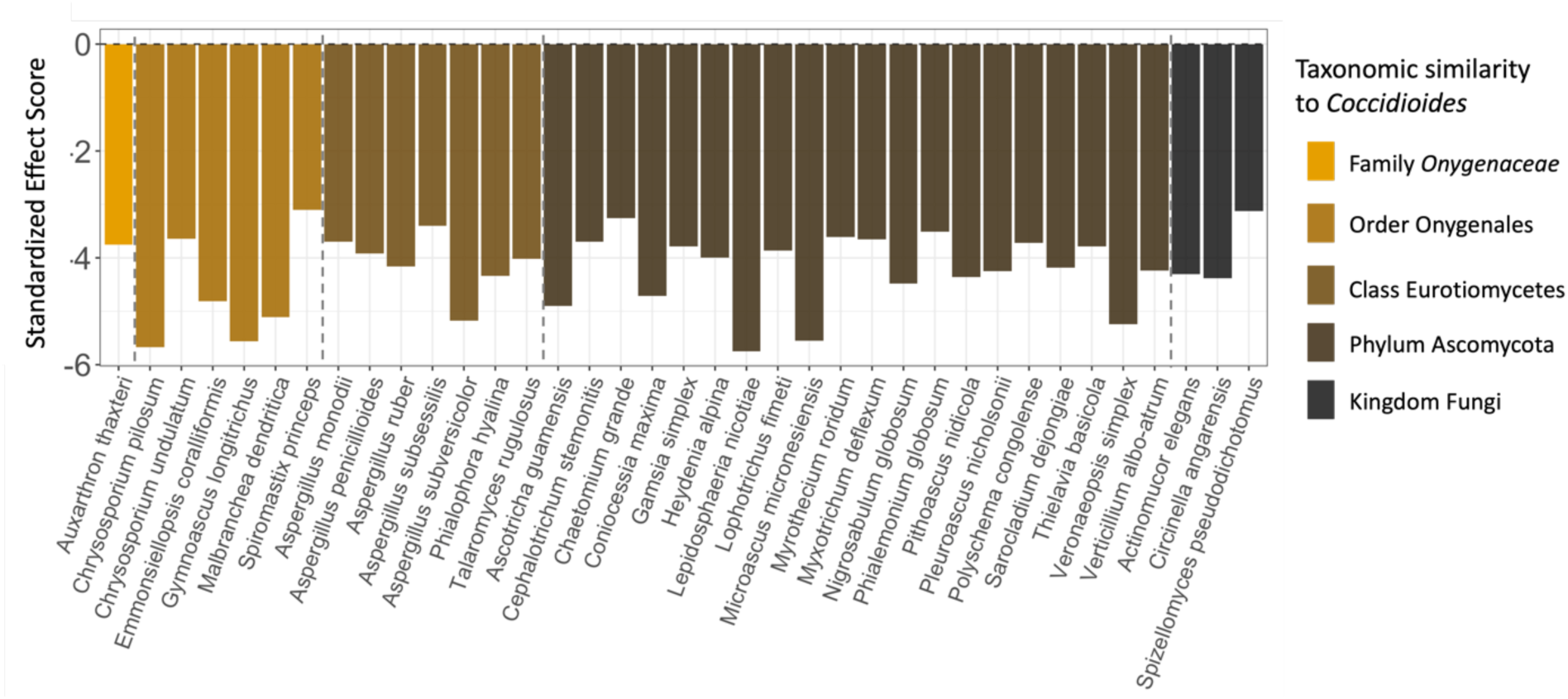
The 37 fungal species that were found to significantly positively co-occur with *Coccidioides* (x-axis), as well as their standardized effect scores (y-axis). Species are color-coded and ordered based on taxonomic similarity to *Coccidioides*, with dashed lines separating the fungal species in the same family, order, class, and phylum as *Coccidioides*. The fungal species that are most similar are on the far left of the plot and least similar on the far right.

## Discussion

Our study identified several key patterns in soil microbial community dynamics of relevance to *Coccidioides immitis* presence in the environment. We found that the largest driver of fungal and bacterial community composition and diversity was the soil microhabitat – namely, whether the soils were derived from within rodent burrows or from the ground surface. That is, microbial communities differed more between soils from the ground surface and rodent burrows—regardless of whether rodents were present—than any other axis of comparison (*i.e.*, rodent exclosure status, *Coccidioides* status). Notably, rodent burrow samples had consistently higher fungal and bacterial diversity than surface soil samples; for fungal diversity, this held true both when rodents were present and excluded. We also found that *Coccidioides*-positive soil samples had higher microbial diversity, particularly fungal diversity, than negative soil samples. Further, *Coccidioides* was found at higher rates in rodent burrow samples, compared to surface soils, as reported elsewhere (14,19,21). Finally, despite prior laboratory evidence demonstrating antagonistic interactions between *Coccidioides* and other soil microbes, we found that no fungal or bacterial species showed negative co-occurrence patterns with *Coccidioides*, while 37 fungal species showed positive co-occurrence patterns.

### The soil microhabitat drives patterns in microbial diversity

There may be several explanations for the differences in microbial diversity between rodent burrows (with or without rodents present) and surface soils. Previous work has shown that rodents affect the soil microbial community through structural creation of burrows (53). Our study site, the Carrizo Plain National Monument, is home to the endangered *D. ingens* (giant kangaroo rat), an ecosystem engineer whose prolific burrow creation is associated with increased plant productivity, invertebrate diversity, and abundance of lizards and squirrels (30). Burrow creation may also play an important role in establishing habitat suitable for diverse bacteria and fungi (53), particularly in desert ecosystems where harsh winds, high temperature fluctuations, and low water availability limit microbial establishment and growth (54). The burrow structure creates a temperature- and moisture-regulated environment that has a more porous soil structure, maintaining favorable conditions for microbes (55–57), including *Coccidioides* (58,59). Our analysis adds to prior evidence using culture-based techniques that found *Dipodomys* spp. burrows host a more diverse fungal community than the surrounding surface soil (53).

We found that soil source (*i.e.*, burrow versus surface soil) was a more important driver of microbial diversity than rodent presence (Figure 4), suggesting that the burrow microhabitat, regardless of rodent activity, provides a favorable environment for diverse microbial taxa in xeric ecosystems. However, we also found that soils from active rodent burrows contained more fungal diversity than soils from inactive burrows, highlighting the important role that rodents play in the ecosystem beyond structural alteration of soil (Figure 2). Rodents may additionally affect the soil microbial community via food caching, quarantining their dead, and urination (59,60). Rodents in the genus *Dipodomys* are known to utilize their burrows and tunnels for various purposes, including nest creation, food caching, temporary shelter construction, and water drainage (61,62), suggesting that burrow entrances may have variable nutrient availability for microbial life based on rodent usage history. Rodent activity may also affect the soil microbial community by interrupting nitrogen uptake by plants and root-associated microbes, potentially increasing nitrogen availability for other microbes in the soil (63). Further, it is possible that urea excreted in rodent urine contributes to *Coccidioides* growth in soils. In its host-associated form, *Coccidioides* has been demonstrated to produce urease to break down urea into ammonia, increasing the alkalinity of its environment and promoting the pathogen’s further growth and virulence (64).

### Coccidioides presence is associated with higher microbial diversity in soils

Our results suggest that the presence of *Coccidioides immitis* in the soil is associated with a more diverse soil microbial community, as the number of fungal species and bacterial families were both significantly higher in *Coccidioides*-positive soil samples than negative samples. These findings are aligned with a prior analysis of nine soil samples collected in Venezuela that showed greater abundance of *C. posadasii* in the soil was associated with greater fungal alpha diversity (27). Taken together with the increased microbial diversity associated with rodents and burrows, and the 37 fungal species found to significantly co-occur with *Coccidioides*, these findings suggest that favorable microhabitats (e.g. rodent burrows) can facilitate a positive association between soil diversity and *Coccidioides* presence. As the association between *Coccidioides* presence and higher microbial diversity remained even after controlling for soil source and rodent presence, other factors unexplored in this study also likely play a role in determining this relationship.

Within the 37 fungal species that significantly co-occurred with *Coccidioides*, those with taxonomic similarity to the pathogen (*e.g.,* same family or order) were overrepresented (Figure 5). This result was surprising as more similar taxa may be expected to occupy similar niches and therefore exhibit greater competition (65). Evolutionary genomics studies have found that taxa in the Onygenales order have lost the ability to bind cellulose, a plant-based nutrient source (66). Further, taxa in the *Onygenaceae* family (including *Coccidioides* spp.) have evolved a shared preference for animal-based proteins such as keratin over plant-based nutrient sources, and therefore may compete for resources in rodent-associated habitats (66–68). Since these taxa occurred together more frequently than expected in this setting, it may be that nutrient availability is not a limiting factor inside rodent burrows, thereby minimizing the role of antagonistic microbial interactions in shaping *Coccidioides* presence in this environment. In addition, rodent burrows may create microclimate conditions to which *Coccidioides* and related taxa are well adapted. Prior work has found that high-temperature, high-salinity conditions in the laboratory stimulate *Coccidioides* growth while inhibiting the growth of microbial antagonists, suggesting that the pathogen may experience fewer competitive interactions in its arid and high-salinity natural environment (28). Habitat filtering, leading to positive associations between phylogenetically similar taxa based on environmental conditions, has been observed as a driver of microbial community composition in other systems, as well (69,70).

Our finding of positive associations between *Coccidioides* and related fungal species, and an absence of antagonistic associations with any fungal or bacterial taxa, contrasts prior findings from laboratory experiments. In particular, *in vitro* co-culture challenge assays of *Coccidioides* with bacteria and fungi isolated from soil at sites known to test positive for the pathogen’s DNA have found microbial antagonists can inhibit the growth of the fungus in the soil through the secretion of antifungal metabolites (25,26). For instance, several *Streptomyces* spp. and *Bacillus* spp. bacterial strains isolated from environmental soil samples near Bakersfield, California exhibited antifungal properties towards *C. immitis* and its close, nonpathogenic relative *Uncinocarpus reesii* in laboratory settings (25). *Bacillus pumilus* and *B. subtilis* were also identified as inhibitors of *Coccidioides* after isolation from soils collected in Arizona, alongside two fungal genera, *Fennellomyces* spp. and *Ovatospora* spp. (26). *Ovatospora unipora* and members of the *Streptomycetaceae* and *Bacillaceae* families were present in our datasets but were not associated with *Coccidioides* presence or absence. Further, our co-occurrence analysis did not identify any species negatively associated with *Coccidioides*, suggesting these antagonistic interactions may be mitigated in *Coccidioides*’ natural environment.

Collectively, these findings suggest that the microbial co-occurrence patterns with *Coccidioides* observed here, along with trends in microbial diversity, are largely influenced by the rodent burrow microhabitat. However, we caution that the microbial associations observed here are not causally interpretable, and it is feasible that other factors are driving the observed microbial co-occurrence patterns.

### Future directions and conclusions

Our amplicon sequencing-based investigation of soil microbial community dynamics in natural settings enabled a more thorough characterization of the overall community than older techniques such as laboratory culture, as many more species are captured through this method (71). However, sequencing may still bias representation toward some taxa over others, and relative abundance based on library preparation does not equate to true abundance in the environment (72,73). Future studies characterizing the abundance and viability of *Coccidioides* in the soil across seasons, interannual climate trends, nutrient availability, and soil types would help resolve the roles of abiotic versus biotic drivers of soil microbial community dynamics.

Our study provides evidence that the soil microhabitat is critical for determining the relationship between the soil microbial community and *Coccidioides* presence. In regions that are highly endemic for the pathogen, small burrowing mammals (including the ecosystem engineer, *D. ingens*) may play a crucial role in altering soil conditions and providing favorable microenvironments for diverse fungal and bacterial taxa, including *Coccidioides* spp. and its relatives in the Onygenales order. Further, we found that the soil microbial community is more diverse in *Coccidioides*-positive soils even when controlling for other variables, suggesting that additional unmeasured factors may play a role in determining this association. While laboratory studies provide crucial insight to potential microbial interactions between *Coccidioides* and other soil microbial taxa, including mechanistic evidence of antagonistic or competitive relationships, these interspecific interactions may not be borne out in natural settings, thus highlighting the importance of continued environmental sampling to uncover the factors driving *Coccidioides*’ presence in the environment.

## Acknowledgements

Research reported in this manuscript was supported by the National Institute of Allergy and Infectious Diseases (NIAID) of the National Institutes of Health under award number R01AI148336. The content is solely the responsibility of the authors and does not necessarily represent the official views of the National Institutes of Health. Thank you to the many people who helped with the collection of soil samples in the field, including Grace Campbell, Erika Lee, Philip Collender, Kate Colvin, and others. Figure 1 was created with BioRender.com.

