## Supplemental data for "Characterizing the soil microbial community associated with the fungal pathogen *Coccidioides immitis*"

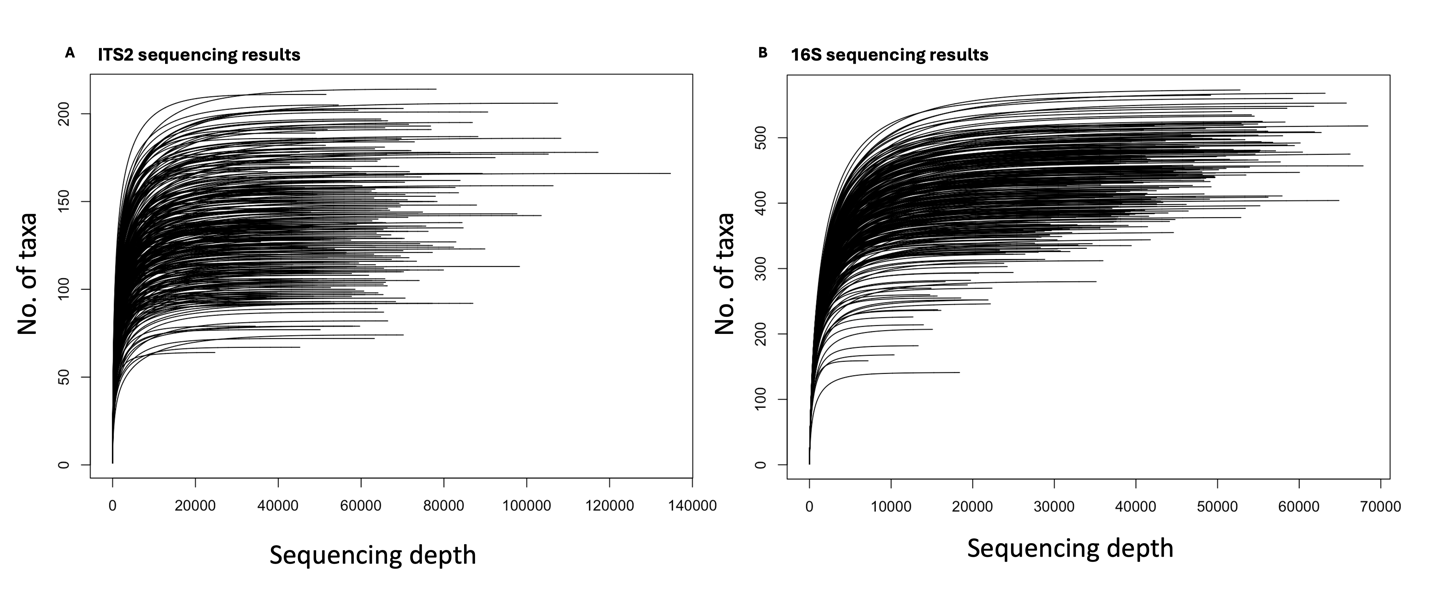

Figure S1. Rarefaction curves for (A) ITS2 and (B) 16S sequencing. Each line denotes a single sequenced sample.

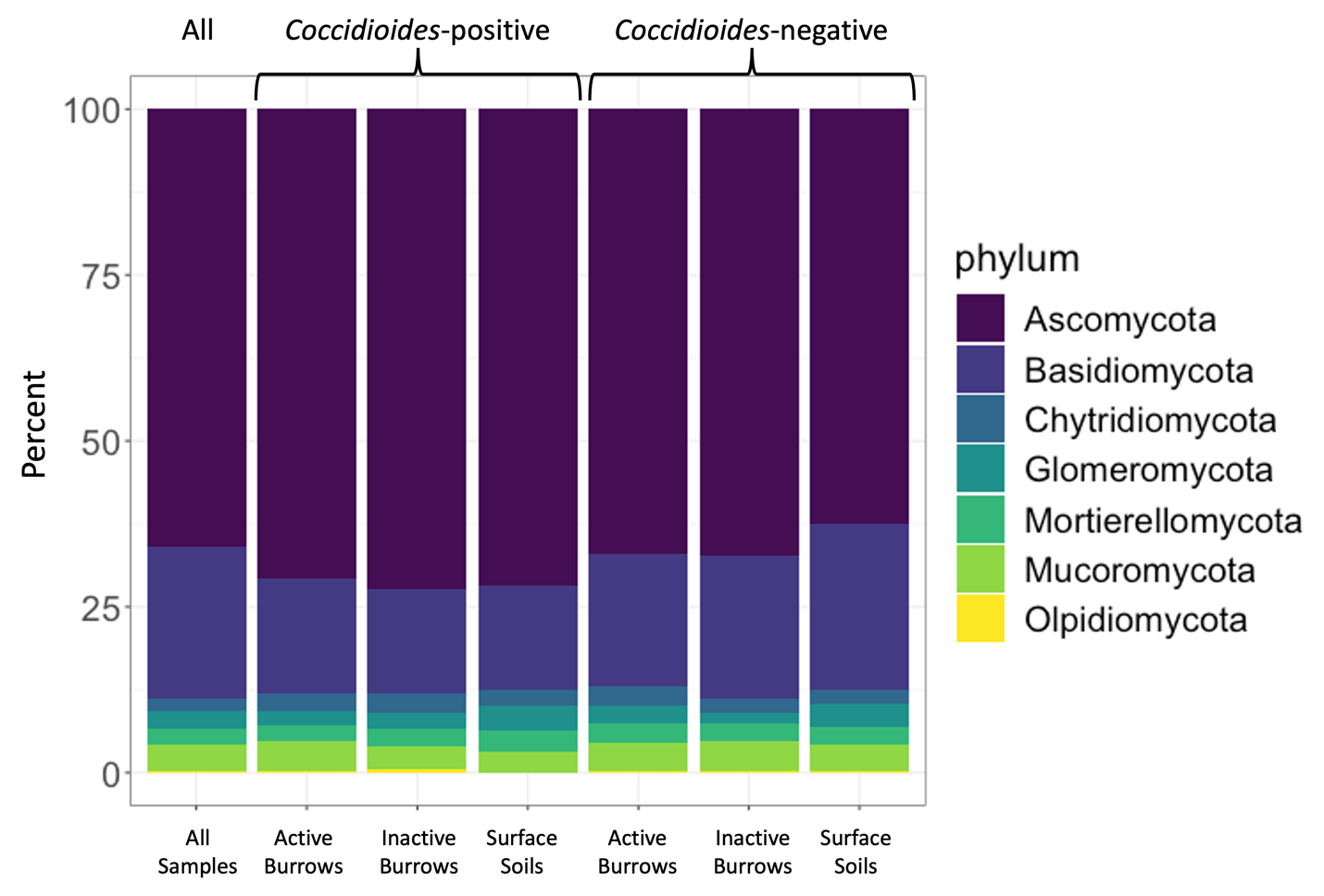

Figure S2. Stacked bar plot showing the proportion of fungal species belonging to each phylum for the full sample set and within sample subgroups. The seven bars are, from the left to right, all samples, *Coccidioides*-positive samples from burrows where rodents are present, from burrows where rodents are absent, and from surface soils; and *Coccidioides*-negative samples from burrows where rodents are present, from burrows where rodents are absent, and from surface soils.

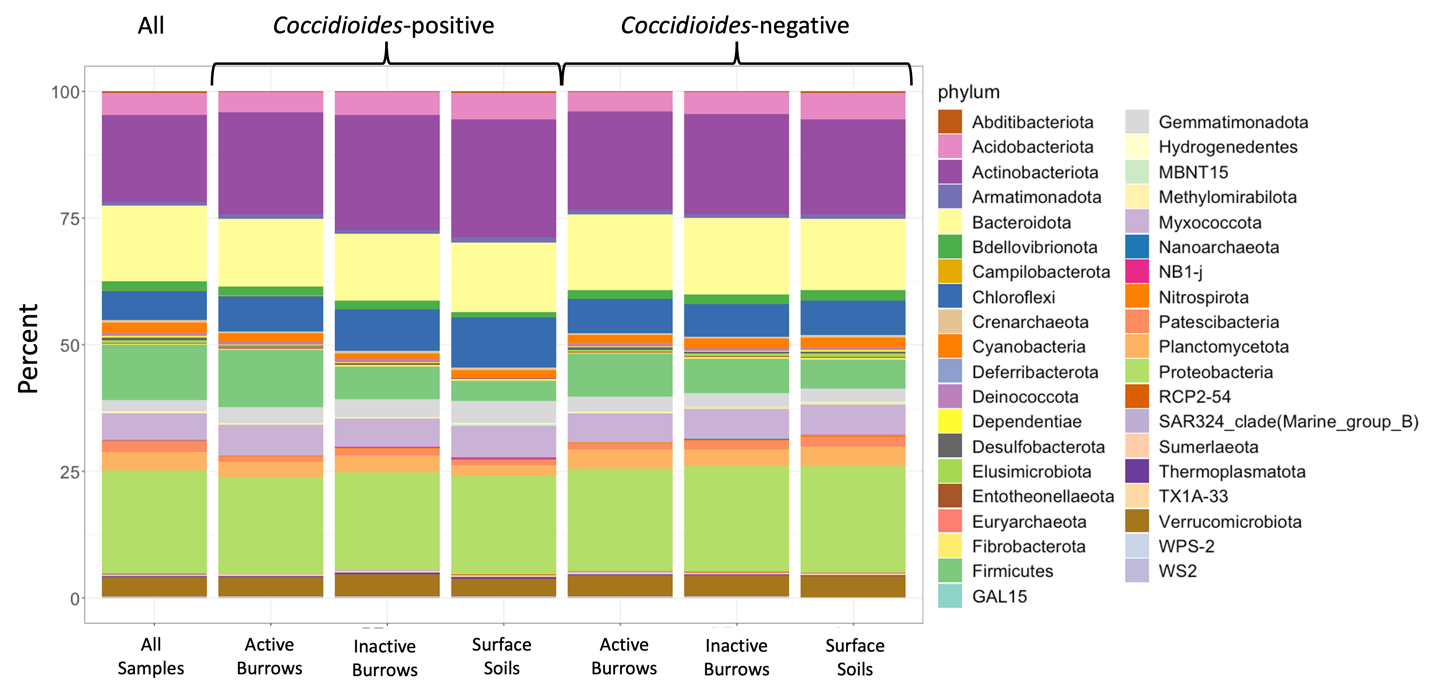

Figure S3. Stacked bar plot showing the proportion of bacterial families belonging to each phylum for the full sample set and within sample subgroups. The seven bars are, from the left to right, all samples, *Coccidioides*-positive samples from burrows where rodents are present, from burrows where rodents are absent, and from surface soils; and *Coccidioides*-negative samples from burrows where rodents are present, from burrows where rodents are absent, and from surface soils.

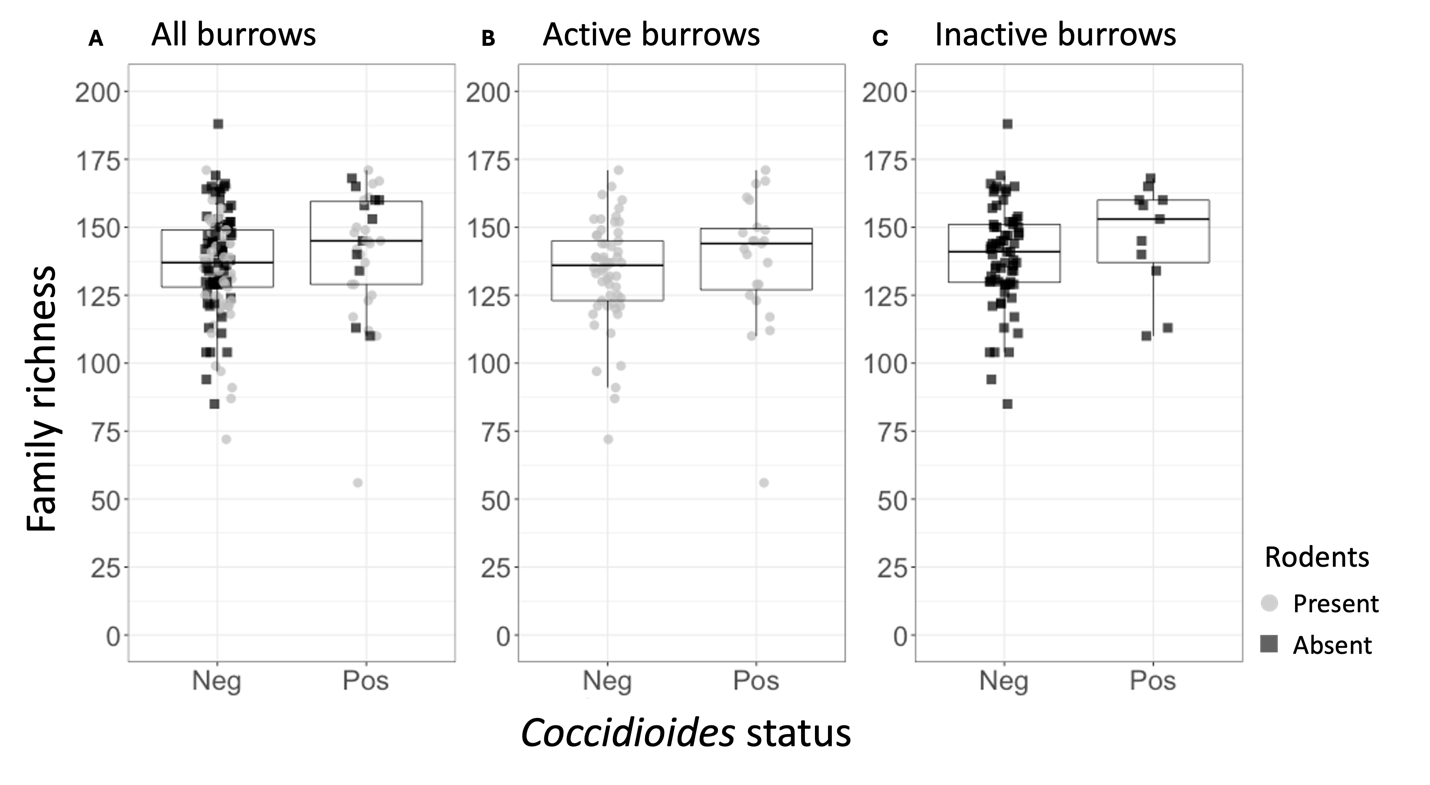

Figure S4. (A) Bacterial family richness for the rodent burrow samples, separated by Coccidioides status (Neg = Coccidioides-negative, Pos = Coccidioides-positive). Points represent individual samples and are color-coded based on whether they were taken from active (gray) or inactive (black) rodent burrows. (B) Bacterial family richness for the active rodent burrow samples, separated by Coccidioides status. (C) Bacterial family richness for the inactive rodent burrow samples, separated by Coccidioides status. Stars indicate degree of significance based on a Wilcox test, such that * = P $\leq$0.05, ** = P $\leq$0.01, *** = P $\leq$0.001, and **** = P $\leq$0.0001. No stars indicate P > 0.05.

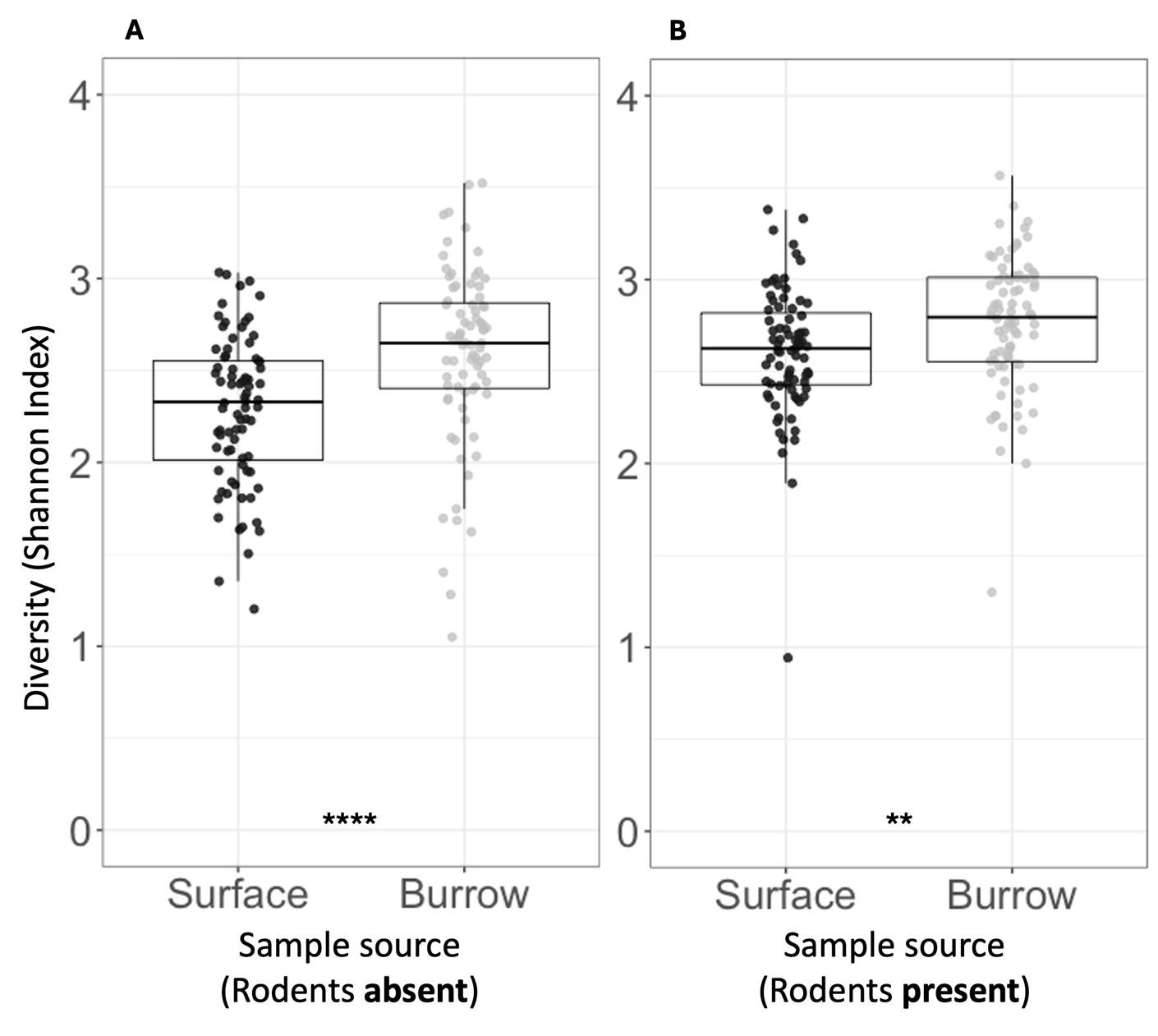

Figure S5. (A) Fungal species diversity for samples taken from the rodent exclosure (where rodents are absent), separated based on soil source (surface or burrow). Points represent individual samples. Samples taken from the ground surface are in black, and those taken from inside rodent burrows are in grey. (B) Fungal species diversity for samples taken from the non-exclosure (where rodents are present), separated based on soil source (surface or burrow). Stars indicate degree of significance, such that * = P $\leq$0.05, ** = P $\leq$0.01, *** = P $\leq$0.001, and **** = P $\leq$0.0001. No stars indicate P > 0.05.

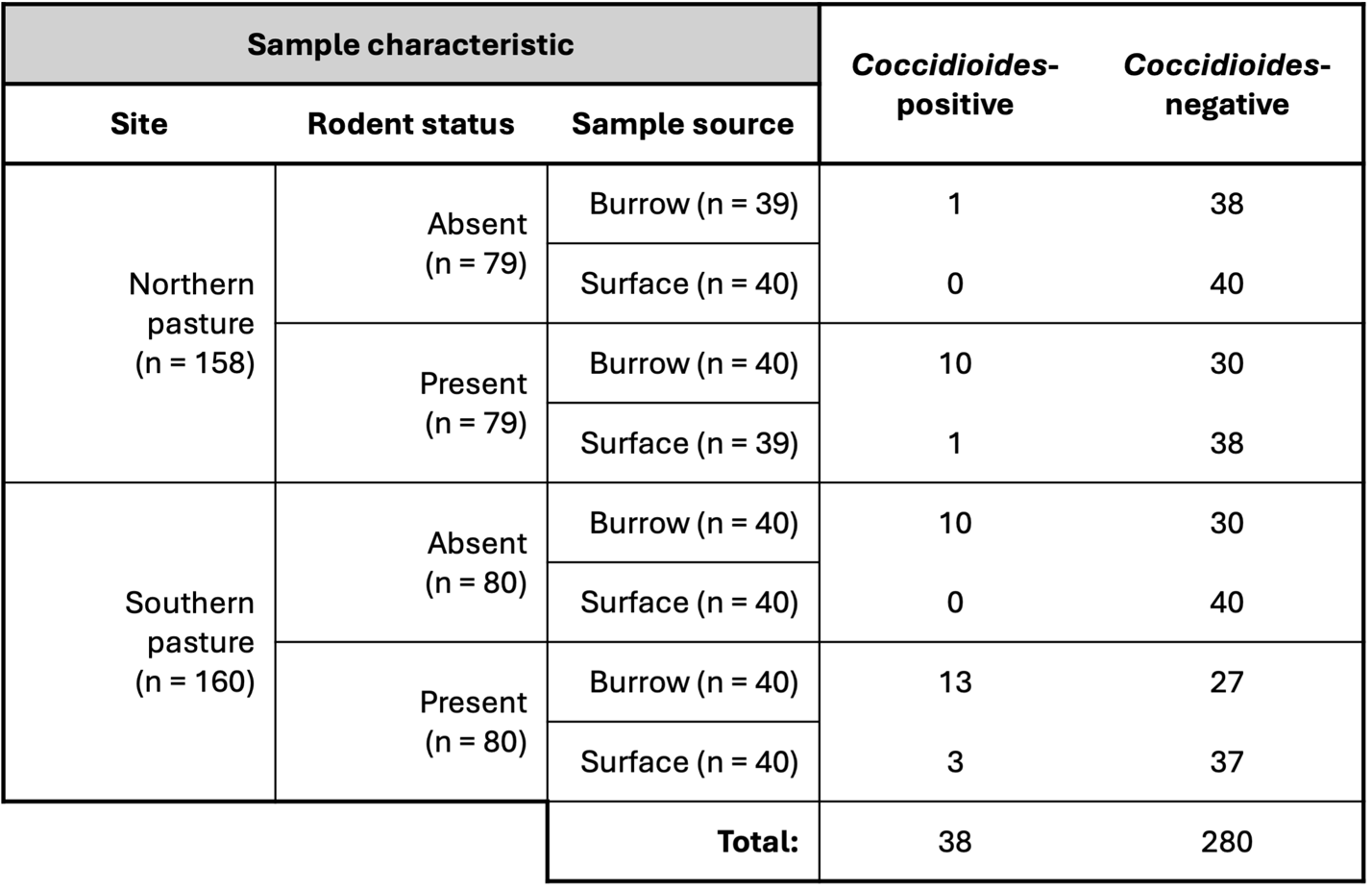

Table S1. Distribution of samples (n = 318) based on sample characteristics, including site, rodent status (rodents absent or present), sample source (burrow or surface), and Coccidioides status (positive or negative).

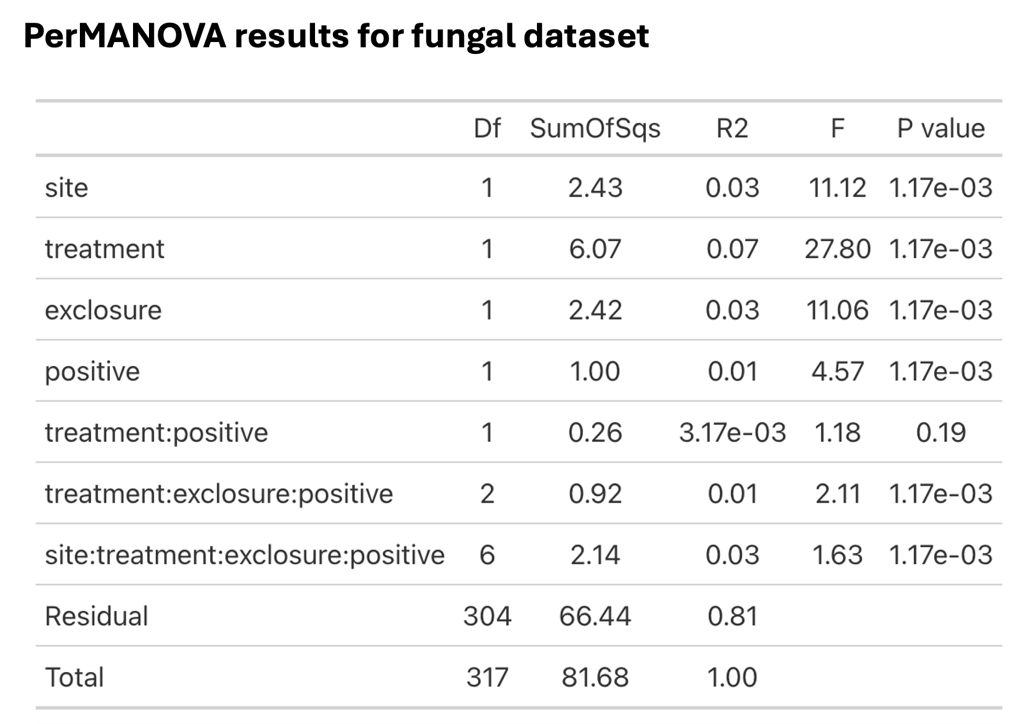

Table S2. Nested PerMANOVA model outputs for the ITS2 dissimilarity matrix, including sum of squares, R^2^, F statistic, and p-values for all individual variables and interaction terms.

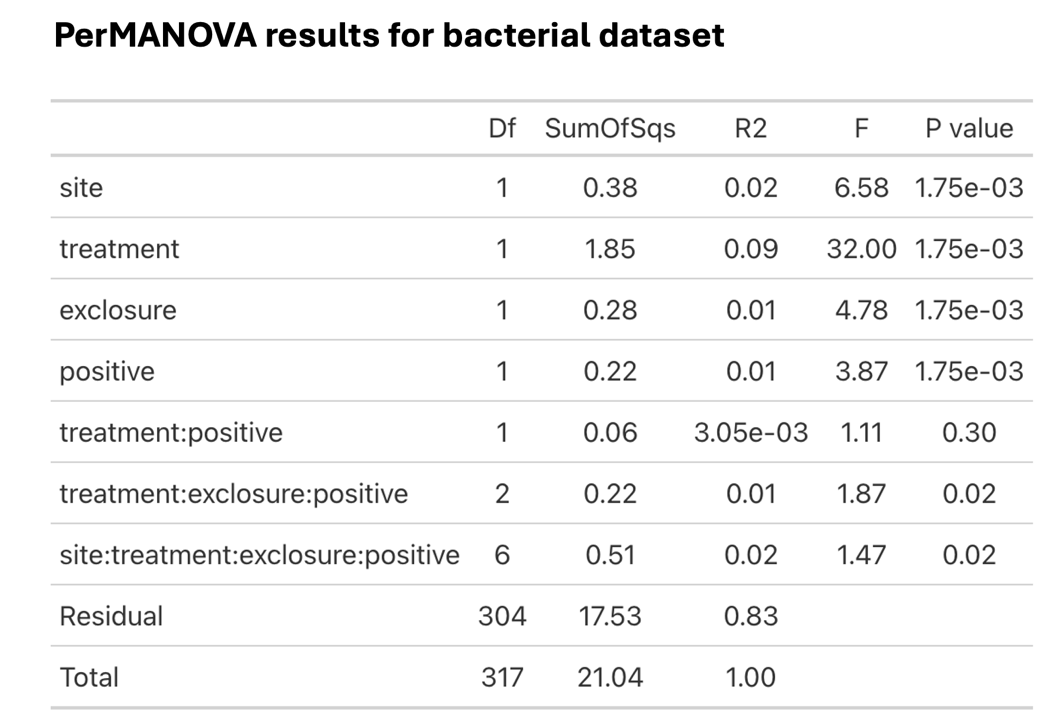

Table S3. Nested PerMANOVA model outputs for the 16S dissimilarity matrix, including sum of squares, R^2^, F statistic, and p-values for all individual variables and interaction terms.
